# Consensus Features Nested Cross-Validation

**DOI:** 10.1101/2019.12.31.891895

**Authors:** Saeid Parvandeh, Hung-Wen Yeh, Martin P. Paulus, Brett A. McKinney

## Abstract

**Motivation:** Feature selection can improve the accuracy of machine learning models, but appropriate steps must be taken to avoid overfitting. Nested cross-validation (nCV) is a common approach that chooses the classification model and features to represent a given outer fold based on features that give the maximum inner-fold accuracy. Differential privacy is a related technique to avoid overfitting that uses a privacy preserving noise mechanism to identify features that are stable between training and holdout sets.

**Methods:** We develop consensus nested CV (cnCV) that combines the idea of feature stability from differential privacy with nested CV. Feature selection is applied in each inner fold and the consensus of top features across folds is a used as a measure of feature stability or reliability instead of classification accuracy, which is used in standard nCV. We use simulated data with main effects, correlation, and interactions to compare the classification accuracy and feature selection performance of the new cnCV with standard nCV, Elastic Net optimized by CV, differential privacy, and private Evaporative Cooling (pEC). We also compare these methods using real RNA-Seq data from a study of major depressive disorder.

**Results:** The cnCV method has similar training and validation accuracy to nCV, but cnCV has much shorter run times because it does not construct classifiers in the inner folds. The cnCV method chooses a more parsimonious set of features with fewer false positives than nCV. The cnCV method has similar accuracy to pEC and cnCV selects stable features between folds without the need to specify a privacy threshold. We show that cnCV is an effective and efficient approach for combining feature selection with classification.

**Availability:** Code available at https://github.com/insilico/cncv.

**Supplementary information:**

## 1 Introduction

Classification and feature selection are fundamental and complementary operations in data mining and machine learning. The quality of selected features affects the quality of the classification model and its performance on validation data (data not used to train and test the model). Specifically, incorporating too many irrelevant features in the training model may lead to predictions that do not generalize well to validation data because the bias-variance tradeoff is tilted towards high variance (overfitting). In contrast, excluding important features from the training model may lead to predictions with low accuracy because the bias-variance tradeoff is tilted towards high bias (underfitting).

There are multiple ways to use feature selection with classification to address the bias-variance tradeoff. Wrapper methods train prediction models on subsets of features using a search strategy to find the best set of features and best model (Guyon and Elisseeff 2003). Optimal wrapper search strategies can be computationally intensive and so greedy methods such as forward or backward selection are often used. Embedded methods incorporate feature selection into the modeling algorithm. For example, least absolute shrinkage and selection operator (Lasso) (Tibshirani 1997) and the more general Elastic Net (or glmnet as it is labeled in the R package) (Zou and Hastie 2005), optimize a regression model with penalty terms that shrink regression coefficients as they find the best model. The Elastic Net hyperparameters are tuned by crossvalidation.

Cross-validation is another fundamental operation in machine learning that splits data into training and testing sets to estimate the generalization accuracy of a classifier for a given dataset (Molinaro, Simon, and Pfeiffer 2005; Kohavi 1995). It has been extended in multiple ways to incorporate feature selection and parameter tuning. A few of the ways CV has been implemented include leave-one-out CV (LOOCV) (Stone 1974), k-fold CV (Bengio, Bengio, and Grandvalet 2003), repeated double CV (Filzmoser, Liebmann, and Varmuza 2009; Simon et al. 2003) and nested CV (Varma and Simon 2006; Parvandeh and McKinney 2018).

Nested CV is an effective way to incorporate feature selection and machine learning parameter tuning to train an optimal prediction model. In the standard nested CV approach, data is split into *k* outer folds and then inner folds are created in each outer training set to select features, tune parameters, and train models. Using an inner nest limits the leaking of information between outer folds that feature selection can cause, and consequently the inner nest prevents biased outer-fold CV error estimates for independent data. However, standard nested CV is computationally intense due to the number of classifiers that must be trained in the inner nests (Wetherill et al. 2018; Varoquaux et al. 2017; Parvandeh et al. 2019). In addition, we show that nested CV may choose an excess of irrelevant features, which could affect biological interpretation of models.

Differential privacy was originally developed to provide useful statistical information from a database while preserving the privacy of individuals in the database (Dwork and Roth 2013). Artificial noise is added to a query statistic high enough to prevent leaking of an individual’s group membership but low enough noise to provide useful group statistical information. This concept of differential privacy has been extended to feature selection and classification with the goal – similar to nested CV – to extract useful information about the outcome (class) variable while limiting information leaking between a training and holdout fold (Dwork et al. 2015). For such machine learning problems, noise is added to the holdout statistic (accuracy or feature importance score) such that zero information from the holdout set is revealed when the difference of the mean statistic between training and holdout stays within a stochastic threshold. This stochastic thresholding procedure that protects the holdout set’s privacy with respect to the outcome variable is known as “thresholdout” (TO). For the smaller sample sizes, typical of bioinformatics data, differential privacy does “preserve the privacy” of the holdout set, but there is a risk of overfitting regardless of the choice of threshold (Le et al. 2017). The private evaporative cooling (privateEC) algorithm (Le et al. 2017), based on concepts of statistical physics and differential privacy, was developed to address the bias-variance balance.

A useful way to interpret the privacy-preserving mechanism in the context of machine learning is that the mechanism creates consistency of the feature importance scores and the accuracy between the training and holdout sets. The proposed consensus nested CV (cnCV) uses this idea of consistency between folds to select features without the need to specify a privacy noise parameter (unlike differential privacy) and without the need to train classifiers in inner folds (unlike standard nested CV). Specifically, cnCV selects top features that are in common across inner training folds. The use of the inner folds prevents information leak between outer folds while using information about the importance rank of features. Furthermore, cnCV extends feature consistency/stability to more folds than simply the two (train and holdout folds) used in differential privacy. The cnCV method uses the nested splitting procedure and selects stable or consensus features first across a given set of inner folds and then across outer folds. The proposed cnCV selects features without training classification models, making it more computationally efficient than nCV. We also show that the consensus features selected are more parsimonious than nCV (fewer irrelevant features).

This new method development study is organized as follows. We describe the new cnCV method, how it differs from standard nCV, and we describe the validation strategy. To validate the methods, we first use simulations with main effects, correlation, and network interactions to compare the classification and feature selection performance of cnCV, nCV, privacy methods TO (Dwork et al. 2015) and pEC (Le et al. 2017), and the embedded method glmnet (Zou and Hastie 2005). Although any feature selection and classification algorithm can be used in cnCV, cnCV, or pEC, we use Relief-based feature selection (Le et al. 2017; Kononenko 1994; Le et al. 2019; Urbanowicz et al. 2018; Le, Dawkins, and McKinney 2019) because of its ability to detect interaction effects as well as main effects. We use random forest as the classifier in cnCV, nCV and pEC because of its known performance and robust hyperparameters. We compare the classification accuracy performance of the methods on a real RNA-Seq study of major depressive disorder. For the RNA-Seq data, we compare accuracies and enrichment of selected genes for known mental disorder associations.

## 2 Methods

### 2.1 Standard nested cross-validation

Nested CV can be used for feature selection and parameter tuning to obtain reliable classification accuracy and avoid overfitting (Varma and Simon 2006; Cawley and Talbot 2010; Tsamardinos, Rakhshani, and Lagani 2014). In standard nested CV (nCV), the dataset is split into *k* outer folds and each fold is held out for testing while the remaining *k* – 1 folds are merged and split into inner folds for training (Fig. 1). Each outer training set is further split into inner folds for inner training and testing. Typically, nCV chooses the outer-fold model that minimizes the inner testing error, but we restrict overfitting by choosing the outer-fold model and hyperparameters with the lowest difference between training and testing accuracy across the inner folds (lowest overfitting). The model and hyperparameters with lowest overfitting across the inner folds is chosen as the training outer loop model and tested on the outer-loop test fold. The features with positive ReliefF scores are used to train the inner fold models. We note that our implementation is not limited to Relief feature selection. The final set of nCV features (final feature selection) are the features used in the best nCV model across the tested outer folds.

**Fig. 1.**
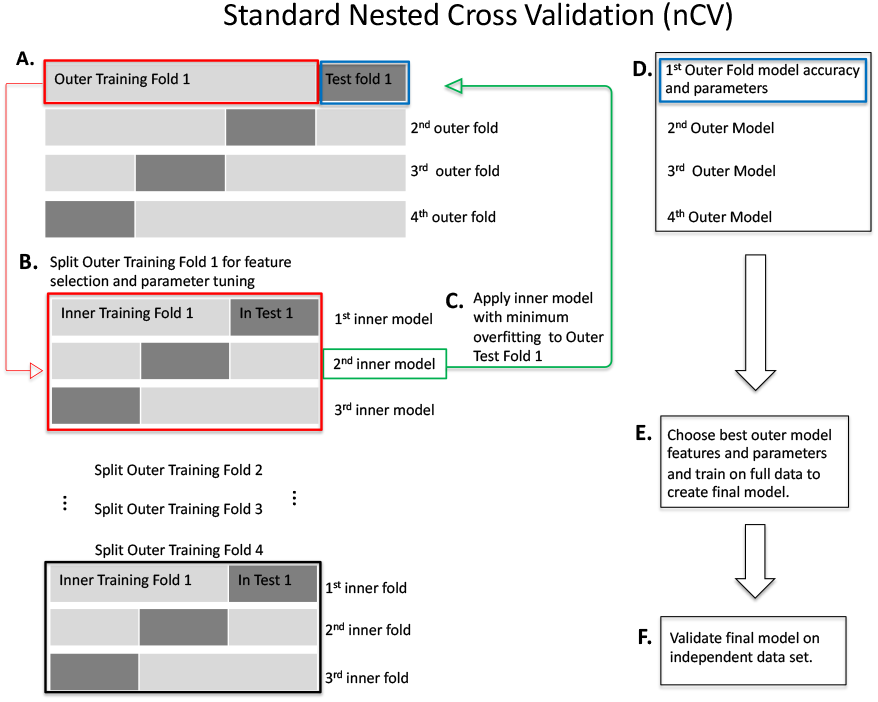
Standard nested Cross-Validation (nCV). **A.** Split the data into outer folds of training and testing data pairs (4 outer folds in this illustration). Then do the following for each outer training fold (illustration starting with Outer Training Fold 1 (red box, A)). **B.** Split outer training fold into inner folds for feature selection and possible hyperparameter tuning by grid search. **C.** Use the best inner training model including features and parameters (2nd inner model, green box, for illustration) based on minimum overfitting (difference between training and test accuracies) in the inner folds to test on the outer test fold (green arrow to blue box, Test Fold 1). **D.** Save the best model for this outer fold including the features and test accuracies. Repeat B-D for the remaining outer folds. **E.** Choose the best outer model with its features based on minimum overfitting. Train on the full data to create the final model. **F.** Validate the final model on independent data.

### 2.2 Consensus nested cross-validation (cnCV)

The cnCV algorithm (Fig. 2) has a similar structure to nCV (Fig. 1), but cnCV has the simplifying feature of not training classifiers in the inner folds. The nCV method selects features and the corresponding model for an outer fold based on the best inner fold classification model. In contrast, cnCV only performs feature selection (not classification) in the inner folds. Features with positive ReliefF scores are assembled for each inner fold and optional parameter tuning is performed (Fig. 2B). Tuning is available in our implementation, but we use standard random forest and ReliefF hyperparameters, where the number neighbors in ReliefF adapts to the number of samples in each fold (Le et al. 2019; Le, Dawkins, and McKinney 2019). The common/consensus features across all inner folds are used to represent the outer-fold set of features (Fig. 2C). Consensus features are chosen again in the outer fold (Fig. 2D) to select the final features for training the final model (Fig. 2E), which is then validated (Fig. 2F).

**Fig. 2.**
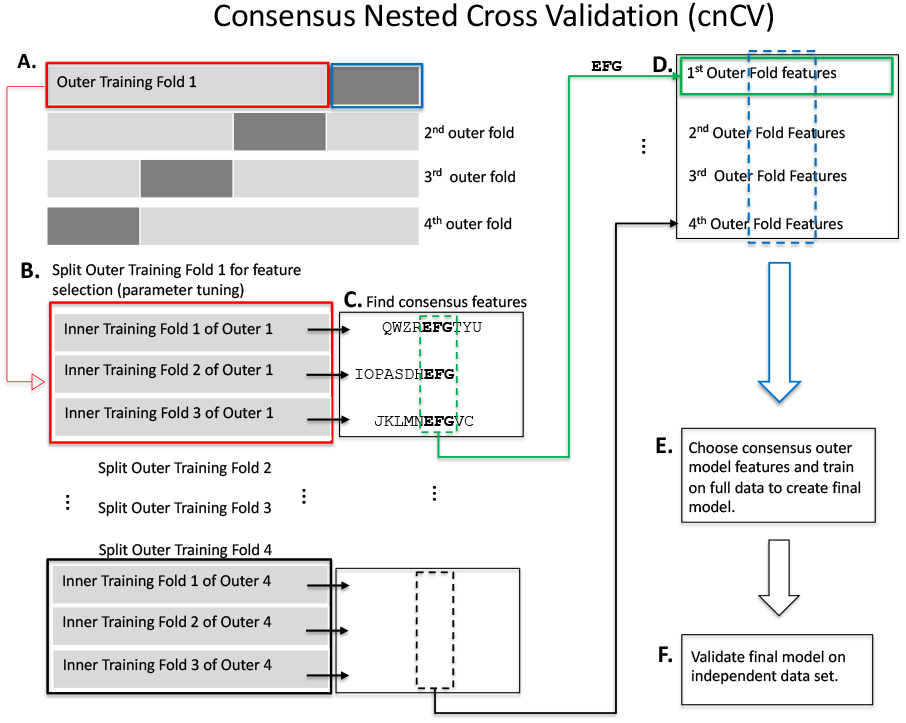
Consensus Nested Cross-Validation (cnCV). **A.** Split the data into outer folds (4 outer folds in this illustration). Then do the following for each outer training fold (illustration starting with Outer Training Fold 1 (red box, A)). **B.** Split outer training fold into inner folds for feature selection and optional hyperparameter tuning by grid search. **C.** Find consensus features. For each fold, features with positive Relief scores are collected (e.g., “QWZREFGTYU” for fold 1). Negative Relief scores have high probability of being irrelevant to classification. The implementation allows for different feature importance methods and tuning the number of input features. Consensus features (found in all folds) are used as the best features in the corresponding outer fold. For example, features “EFG” are shared across the three inner folds. This procedure is used in the inner and outer folds of cnCV. Classification is not needed to select consensus features. **D.** The best outer fold features (green arrow to green box) are found for each fold (i.e., Repeat B-D for all outer folds). **E.** Choose the consensus features across all the outer folds to train the final model on full data. Consensus features are selected based on training data only. Classification is not performed until the outer consensus features are selected (A-D). **F.** Validate the final model on independent data.

For cnCV, nCV and pEC in the current study, we use the nearest-neighbor-based ReliefF feature selection algorithm because of its ability to detect main effects and interactions. For cnCV, we use all positive Relief scores (in Fig. 2B) as a conservative threshold because negative Relief scores are likely irrelevant to classification. Relief scores are the difference of the variable’s values between nearest neighbors from the opposite phenotype group and the same phenotype group, and a negative score means the variable discriminates poorly between groups. Positive scores may include false positives but the consensus procedure helps to eliminate these. If the user chooses another feature selection algorithm, a similar conservative threshold or top percentage of features can be used and then the consensus mechanism will remove features that are unstable across folds. The software implementation includes an option for tuning the threshold.

Our implementation includes multiple Relief neighborhood options, but for the current study we use a fixed number of nearest hit and miss neighbors, *k_R_* = 0.154(*m*’ – 1), that adapts to the number of samples *m*’ in a fold. The 0.154 prefactor yields an approximation to a fixed radius that contains neighbors within a half standard deviation of a sample’s radius in the attribute space (Le et al. 2019; Le, Dawkins, and McKinney 2019). This value for the number of Relief neighbors has been shown to provide a good balance for detecting main effects and interaction effects (Le et al. 2019). We use random forest for classification with 500 trees (ntree) and we used p/3 as the number of random features chosen as candidates for splitting each node (mtry), where p is the number of available features in a fold. The software implementation allows for the tuning of the random forest and Relief parameters, but we fix the values because of the number of simulation analyses and to focus on the comparison of the consensus feature selection. In the thresholdout (TO) method in the current study, we use a univariate feature selection and Naïve Bayes classifier, and we use a threshold noise parameter of 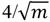, where m is the number of samples.

### 2.3 Simulation approach

We use the privateEC package (https://github.com/insilico/privateEC) to create multiple types of simulated data with main effects and interaction network effects to assess the CV methods (Le et al. 2017). For each replicate dataset, we generate a training set with m=200 samples and a validation set with 100 samples to test true generalizability. Each simulated dataset has balanced cases and controls. We choose a sample size consistent with real gene expression data but on the smaller end to demonstrate a more challenging scenario. We apply each method to a training dataset and store the final set of features and model with its holdout accuracy. The final holdout model from the training set is tested on an independent set for true generalization accuracy.

Each data set contains p=500 variables. We also include results with 10,000 variables (See Supplementary Materials). We create simulations with main effects and interaction effects, where 10% (50) of the attributes are functional or true associations (see Refs. (Le et al. 2017, 2019; Lareau et al. 2015) for details of the simulation method). The simulations are replicated 100 times with noise to compute average performance of methods.

We use three categories of effect size: easy to detect (40% power), medium (30% power) and hard to detect (20% power) (Le et al. 2017). For main effects, we model multiple independent random normal effects. In addition, we vary the strength of correlation between variables. The interaction effects are created from a simulated differential co-expression network, and the effect size is a noise parameter that controls the correlation. This correlation is related to the interaction effect size because we disrupt the correlation of target genes in cases but maintain correlation within controls, thereby creating a final differential correlation (interaction) network.

### 2.4 Software availability

All algorithms and analysis code for reproducibility are available as an R package on github, https://github.com/insilico/cncv.

## 3 Results

### 3.1 Simulation comparison

For the nested CV methods, we used 10 inner and outer CV folds. The classification performance of the comparison methods (cnCV, nCV, TO, pEC and glmnet) is similar for simulated data with main effects (Fig. 3 A-C) and for main effects with correlation (Fig. 3 D-F), but TO and glment overfit more than the other methods. For correlated data, the average correlation is 0.8 for connected variables in the gene network and 0.2 for unconnected variables. In simulated data with interaction effects (Fig. 3 G-I), glmnet and TO are not able to classify the data because they assume additive effects. The cnCV, nCV and pEC are able to classify the interaction data because they use Relief feature selection. The comparison methods show the same trends when we increase the number of attributes from p=500 to 10,000 (See Supplementary Material).

**Fig. 3.**
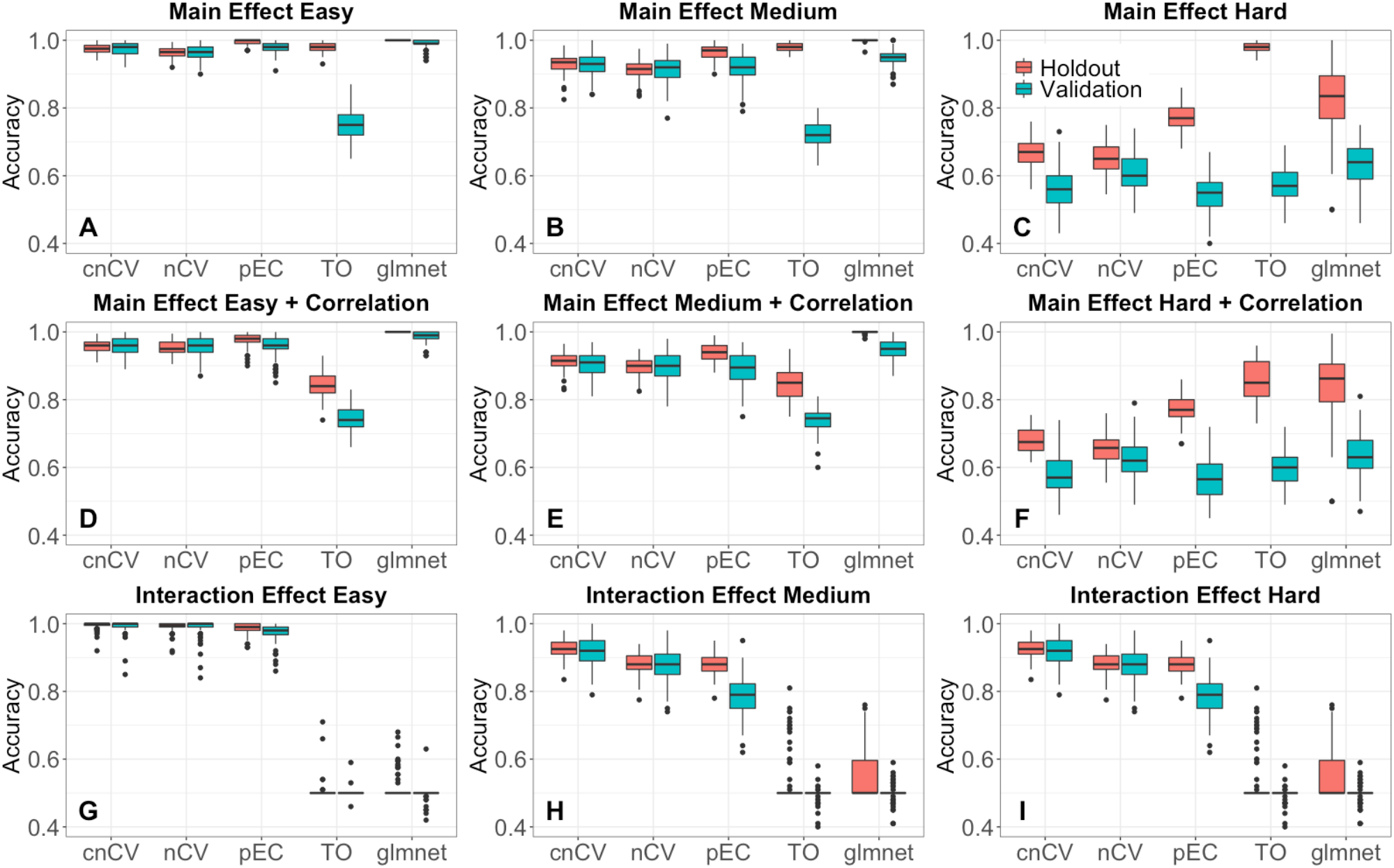
Simulated training/holdout and validation accuracy comparison. Accuracies for consensus nested CV (cnCV), standard nested CV (nCV), private Evaporative Cooling (pEC), differential privacy thresholdout (TO) and glmnet for 100 replicate simulated datasets with main effects (A-C), main effects with correlation (D-F) and interaction effects (G-I). Training/holdout data (red) has m=200 samples, balanced cases and controls, and validation data (teal) has m=100 samples. Effect sizes range from easy to hard (left to right) with p=500 variables, 10% functional effects. Red boxplots indicate holdout accuracies (final holdout model from training) and teal boxplots indicate validation accuracies (final holdout model applied to independent data). Accuracies for all methods (except glmnet) are computed from random forest out-of-bag. Glmnet accuracies are computed from the fitted model coefficients and optimal elastic-net lambda and alpha parameters tuned by the cross-validation.

We also compare the feature selection performance of cnCV, nCV, TO, pEC and glmnet (Fig. 4) using precision and recall to detect the 50 functional simulated features out of 500. Standard nCV has a tendency to include too many features in its models (253 averaged across all simulations) compared to the true number of functional features (50). This leads to nCV returning more false positives and lower precision than the other methods. Despite the large number of false positives, nCV still has high classification accuracy (Fig. 3), which is likely a reflection of the robustness of random forest to irrelevant features when a sufficient number of relevant features is included in the model. For cnCV on the other hand, the number of features selected (43 on average across all simulations) is closer to the true number of functional (50), and cnCV has a higher precision than nCV. Glmnet has good precision for the main effect models (Fig. 4 A-C) but cannot detect interaction effects (Fig. 4 G-I) as expected due to additive model assumptions. In the easy interaction effect simulations (Fig. 4G), TO and glmnet have very wide bars for precision because they select very few variables and, as the interaction is relatively easy, in about half of the simulations they select 1 functional variable and other times none.

**Fig. 4.**
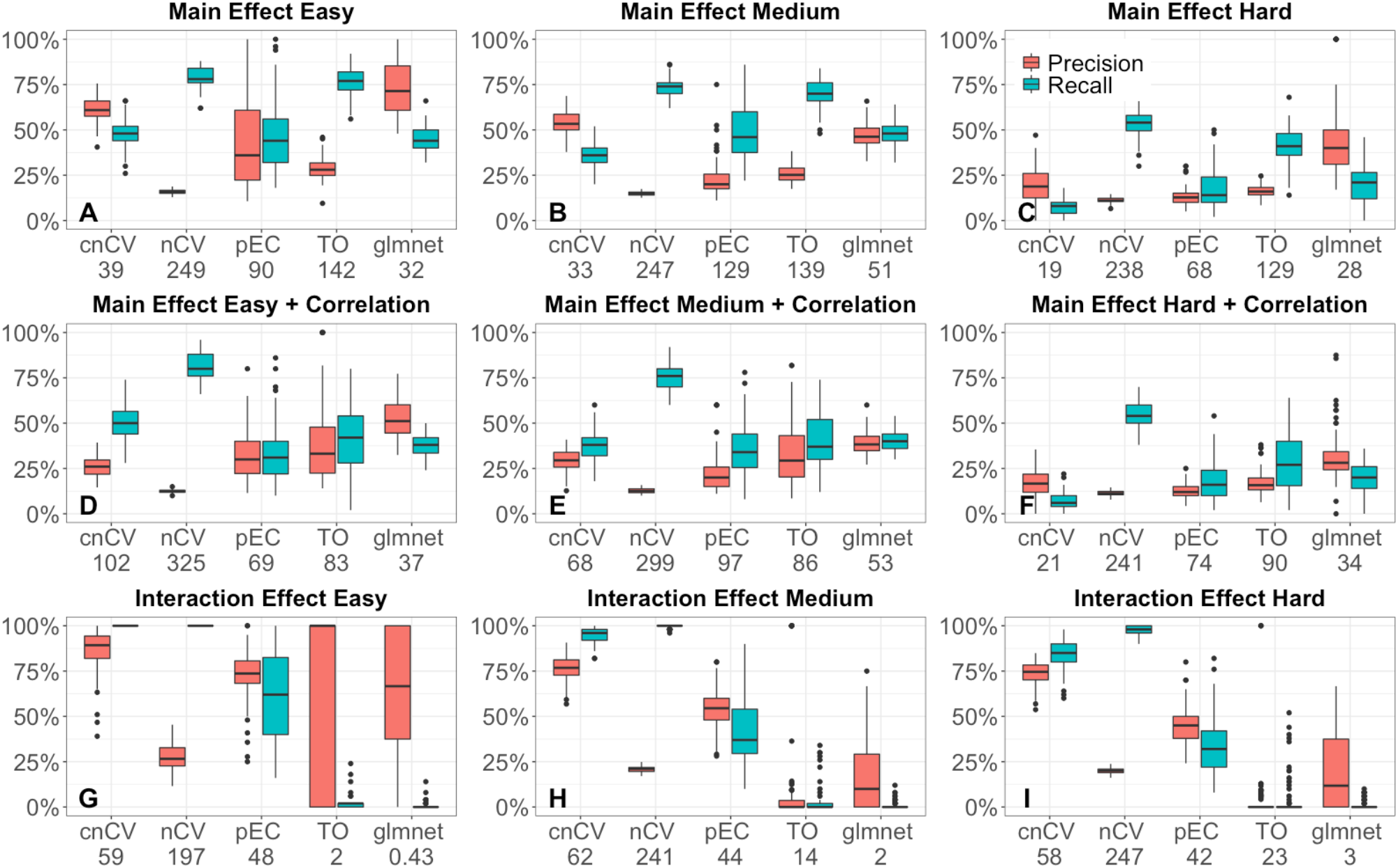
Precision (red) and recall (teal) for selecting functional features (50 functional out of p=500 and m=200 samples) for consensus nested CV (cnCV), standard nested CV (nCV), private Evaporative Cooling (pEC), differential privacy thresholdout (TO) and glmnet for 100 replicate simulated datasets with main effects (A-C), main effects with correlation (D-F) and interaction effects (G-I). Effect sizes range from easy to hard (left to right). The number below each method (on horizonal axes) is the average number of features selected by the method. Precision and recall computed from the training/holdout data.

### 3.2 Analysis of RNA-Seq data

In addition to simulated data, we compare classification accuracy using real RNA-Seq data for MDD (Mostafavi et al. 2014), with 15,231 genes and 915 subjects with 463 major depressive disorder and 452 controls (Fig. 5). We filtered 50% of genes using coefficient of variation, which resulted in 7,616 genes for analysis. We train random forest classifiers by cnCV (399 genes selected), nCV (3,592 genes selected), and pEC (3,415 genes selected) with adaptive-neighbor ReliefF feature selection. For glmnet, we optimize the hyperparameter λ (cv.glmnet) with a=1 (Lasso). The glmnet curve of CV error versus λ is very flat near the minimum, and we choose the value of λ near the CV minimum that selects the most features because glmnet tends to remove many features (56 genes selected). We compare the performance of the methods on real data by splitting the samples into a training half (Fig. 5, left) and a validation half (Fig. 5, right) to compare the utility and generalizability of the methods. Glmnet has the highest accuracy on the training data; however, cnCV has the highest accuracy on the validation data. The validation accuracy is a more realistic measure of the methods’ performance. In addition to having a higher validation accuracy than glmnet, the cnCV training accuracy is much closer to its validation accuracy indicating less overfitting. The glmnet classifier has similar overfitting regardless of λ values near the CV error minimum. The nCV model has similar low overfitting to cnCV but its accuracies are lower than cnCV.

**Fig. 5.**
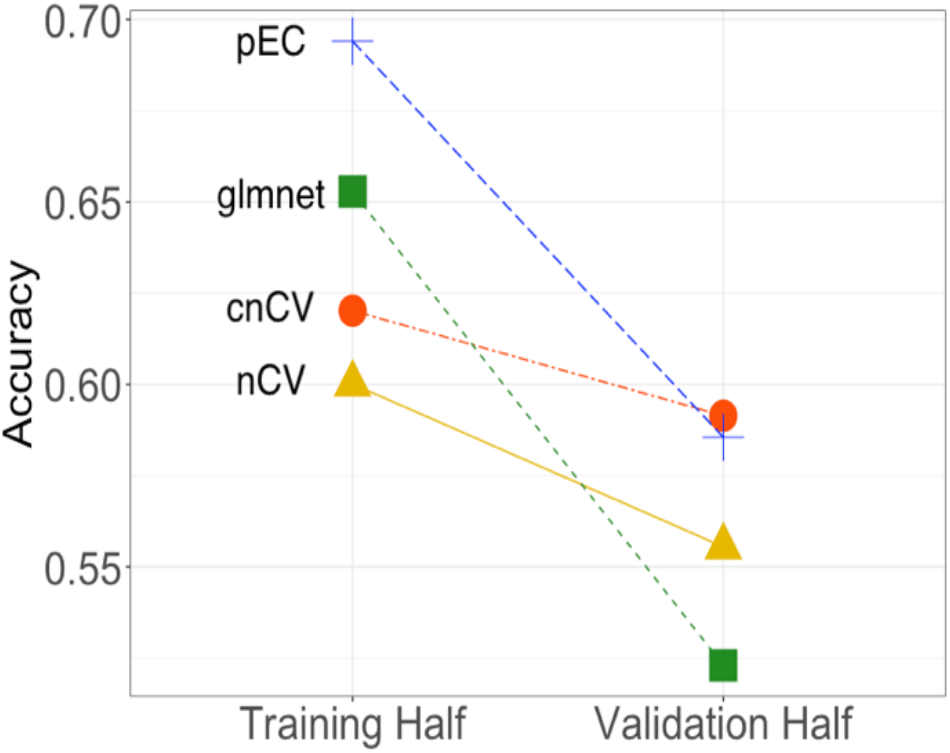
RNA-Seq accuracy comparisons. RNA-Seq dataset from a major depressive disorder study (Mostafavi et al. 2014) is split into a Training Half and a Validation Half. The final training model for each method is tested on the independent validation half to assess the degree of overfitting. Standard nested CV (nCV, triangle) uses 3,592 genes in its model, consensus nested CV (cnCV, circle) uses 399 genes in its model, and private Evaporative Cooling (pEC, plus) uses 3,415 genes in its model. These methods use ReliefF feature selection and random forest classification. Glmnet (square) selects 56 genes by penalized logistic regression feature selection. The cnCV validation accuracy is highest and has the lowest overfitting.

**Table 1.** Runtimes averaged over 100 simulated datasets with m=200 samples and p=500 attributes (second and third columns) and m=200 samples and p=10,000 attributes (right two columns) using a 3.1 GHz Intel Core i7 CPU with 16GB of RAM.

| Method | 500 attributes |  | 10,000 attributes |  |
| --- | --- | --- | --- | --- |
|  | 200 samples |  | 200 samples |  |
|  | Mean | Standard | Mean | Standard |
|  | (seconds) | Deviation | (seconds) | Deviation |
| cnCV | 16.55 | 0.91 | 1,756.54 | 66.11 |
| nCV | 1,88.92 | 127.84 | 40,607.38 | 7,380.84 |
| pEC | 28.09 | 0.77 | 21,866.94 | 6,969.38 |
| TO | 0.04 | 0.02 | 6.13 | 0.05 |
| glmnet | 0.48 | 0.12 | 6.08 | 0.72 |

We use DisGeNET to assess the disease relevance of genes that were selected by each method out of the initial 7,616 filtered genes. Dis-GeNET is a curated repository of collections of genes and variants associated with human diseases from GWAS catalogs, animal models and the scientific literature (Piñero et al. 2017). We queried the repository for the category “Mental or Behavioral Dysfunction” with genes selected by cnCV (399), nCV (3,592), pEC (3,415) and glment (56). See Supplementary Material for list of overlapping genes. The overlap with the dysfunction category is partly driven by the number of genes selected. The two methods that included the most genes had the largest overlap with the DisGeNET category: nCV (959 overlap out of 3,592, P=7e-33) and pEC (925 overlap out of 3,415, P=2e-28). The methods that selected fewer genes had lower overlap and lower enrichment significance: cnCV (116 overlap out of 399, P=.004) and glmnet (17 out of 56 overlap, P=.19).

## 4 Discussion

Nested cross-validation incorporates feature selection and parameter tuning into machine learning model optimization to improve model accuracy while limiting the effects of overfitting. Nested cross-validation can be computationally intense and it can select many extraneous features. We developed a consensus nested cross-validation (cnCV) method that improves the speed of nested CV and selects a more parsimonious set of features. Speed is improved by choosing consensus features across inner folds instead of using classification accuracy to select features and models.

The consensus selection approach is motivated by the objective of finding consistent or stable features across folds, which is related to the thresholdout (TO) mechanism of the differential privacy-based framework. The TO mechanism prevents leaking of information between two folds (training and holdout) by adding noise to holdout-fold scores. Unlike TO differential privacy, cnCV is effective for lower sample sizes and can use any number of CV folds. The cnCV approach also does not require a privacy noise threshold, although it does use a fixed number or threshold of top input features.

In the consensus method, we included all ReliefF features with nonnegative scores because negative Relief scores are unlikely to be useful for classification. We hypothesize that the features selected by cnCV are similar to using a multiple-testing correction of P values from a statistical inference Relief (Le et al. 2019) or a projected distance regression (Le, Dawkins, and McKinney 2019). For other feature selection methods, one may wish to use a threshold number of top features or a significance threshold in cnCV. Choosing a threshold that allows too many features may increase overfitting because of the increased risk of finding features that overlap by chance. However, if false features are selected, the outer test fold should reflect this and give a correct (lower) generalization accuracy. On the other hand, choosing a threshold that allows too few features may reduce the accuracy of the classifier. The number of input features may be addressed by tuning the threshold in the inner folds, which is implemented in the software.

Our main goal was to compare standard nested CV with the new cnCV. Thus, for both we used the same feature selection (ReliefF) and classification method (random forest). We used ReliefF as the feature selection algorithm because of its ability to detect main effects and interactions. The cnCV implementation can also be used with NPDR to adjust for covariates such as sex (Le, Dawkins, and McKinney 2019). Although it uses a different feature selection and classification approach, we also compared with glmnet because of its wide representation as a method that uses CV for model optimization. Our cnCV implementation is not limited to Relief feature selection or random forests for classification and it applies to regression problems (continuous outcomes) and includes parameter tuning.

