## Supplemental Figure 1 for "Consensus Features Nested Cross-Validation"

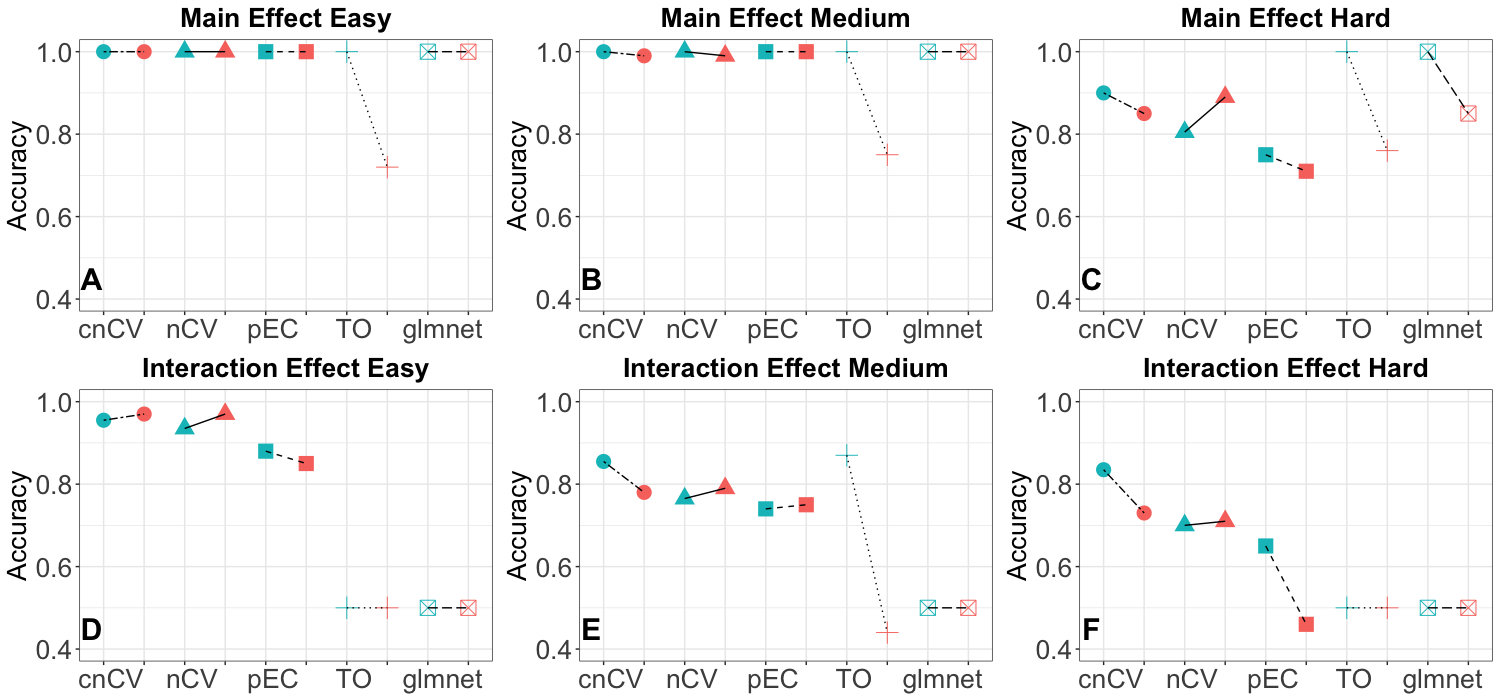

**Figure S1.** **Average accuracies 10,000 attributes**. Training/holdout (teal) and validation (red) accuracies for consensus nested CV (cnCV), standard nested CV (nCV), private Evaporative Cooling (pEC), differential privacy thresholdout (TO) and glmnet for 10 replicate simulated datasets with main effects (A-C), main effects with correlation (D-F) and interaction effects (G-I). Training data has m=200 samples, balanced cases and controls, and validation data has m=100 samples. Effect sizes range from easy to hard (left to right) with p=10,000 variables, 10% functional. Pairs of plot points indicate holdout accuracies (final holdout model from training, teal) and validation accuracies (final holdout model applied to independent data, red) for each method. Accuracies for all methods (except glmnet) are from random forest out-of-bag. Glmnet accuracies are computed from the fitted model coefficients and optimal elastic-net lambda and alpha parameters tuned by the cross-validation.

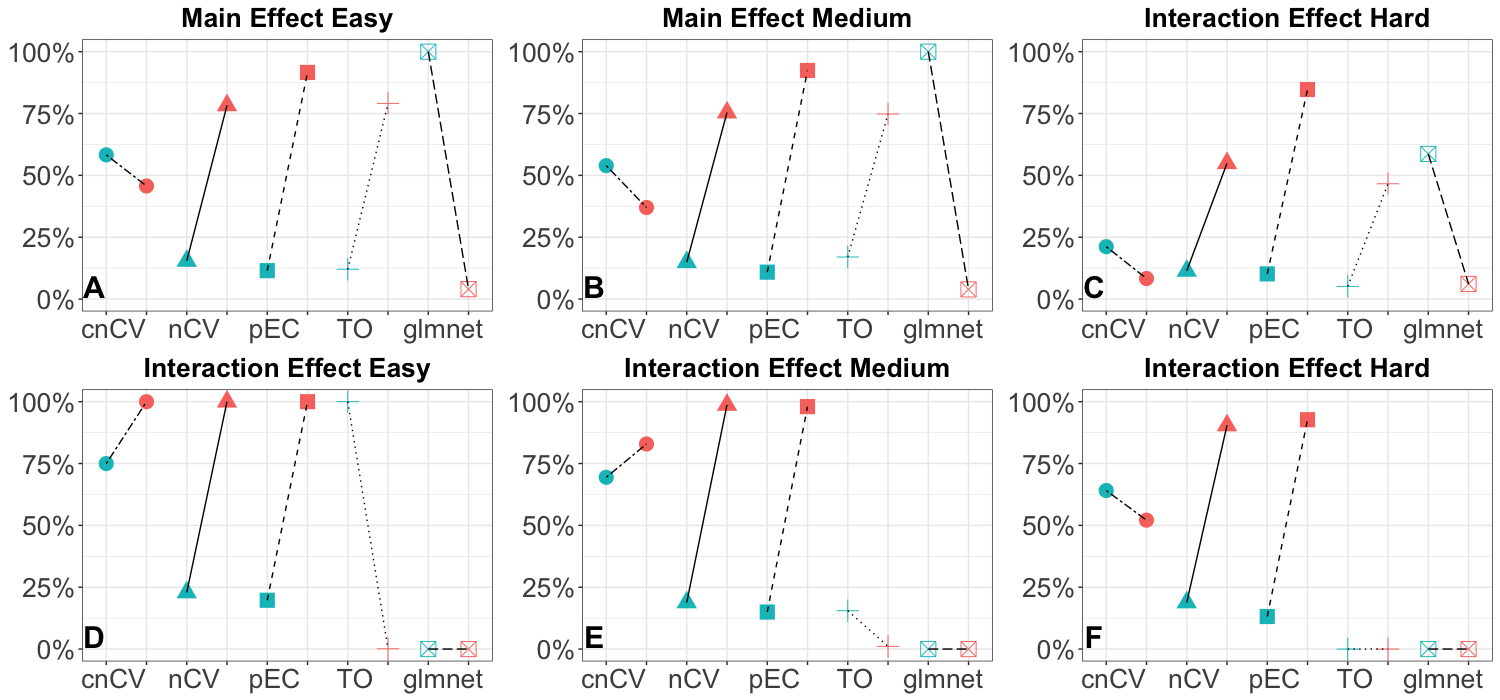

**Figure S2. Precision and recall for 10,000 attributes**. Comparison of precision (teal) and recall (red) for selection of simulated functional features (10% functional out of p=10,000 and m=200 samples) for consensus nested CV (cnCV), standard nested CV (nCV), private Evaporative Cooling (pEC), differential privacy thresholdout (TO) and glmnet for 10 replicate simulated datasets with main effects (A-C), main effects with correlation (D-F) and interaction effects (G-I). Effect sizes range from easy to hard (left to right).
